# Achilles’ Heel: Type VII Toxin/Antitoxin System Hha/TomB Facilitates Phage Infection by Regulating Cryptic Prophage Lytic Genes

**DOI:** 10.64898/2026.10.29.762910

**Authors:** Eunsol Seo, Maura E. Cingle, Hadar Ben-Yoav, Ilana Kolodkin-Gal, Thomas K. Wood

**Author notes:** For correspondence.(+)1.

## Abstract

Toxin/antitoxin systems (TAs) are stress-response loci most-closely linked to stabilizing mobile genetic elements such as plasmids and inhibiting phage infection. In contrast to the phage inhibition paradigm, here we demonstrate that the type VII TA system, Hha/TomB, *facilitates* rather than *inhibits Escherichia coli* infection with 12 BASEL lytic phages (seven *Tequatrovirus* phages and five *Vequintavirus* members), including phage 48 (a *Vequintavirus*), which we chose to focus on due to its clear phenotype. Toxin Hha regulates cryptic prophage lytic genes, and antitoxin TomB masks Hha toxin activity by oxidizing a conserved Hha cysteine residue. We find here that the mechanism by which Hha increases infection is via activation of cryptic prophage lytic genes. Therefore, this is one of the first reports of phages and a type VII TA system as well as the first report of a TA system that is not advantageous for phage inhibition but instead may be an Achilles’ heel that is manipulated by lytic phages.

## INTRODUCTION

Toxin/antitoxin systems (TAs) are ubiquitous stress-response loci most-commonly linked to mobile element stabilization and phage defense (Song and Wood, 2020). TAs are classified into nine types (Seo et al.) based primarily on how the antitoxin neutralizes the toxin, which slows growth by corrupting targets such as DNA replication, cell division, mRNAs/tRNAs, and energetic molecules such at ATP (Jurėnas et al., 2022). The first TAs found to inhibit phage was the type I system Hok/Sok in 1996 (Pecota and Wood, 1996); since then, myriad types of TAs have been shown to defend the cell from phages including type II MazE/MazF (Hazan and Engelberg-Kulka, 2004), type III ToxN/ToxI (Fineran et al., 2009), type IV ShosT/ShosA (Pu et al., 2025), type VII PfkAB/PfpC (Guo et al., 2024), and type IX CloA/CloB (Clover) (Yamaguchi et al., 2025). However, there is no report of the founding type VII TAs Hha/TomB (Wang et al., 2021) and phage inhibition.

In type VII TAs (Wang et al., 2021), the antitoxin directly inactivates the toxin as an enzyme. For example, antitoxin TomB (toxin <u>o</u>verexpression <u>m</u>odulator in <u>b</u>iofilms) oxidizes Cys18 of toxin Hha (<u>h</u>igh <u>h</u>emolysin <u>a</u>ctivity) to inactivate it (Marimon et al., 2016). Furthermore, toxin Hha is induced in biofilms (Ren et al., 2004) and functions with protein partner H-NS, a global transcription repressor of foreign DNA (Rimsky, 2004), by binding to AT-rich DNA regions. Specifically, Hha represses biofilm formation by activating cryptic prophage lytic genes by binding to at least 15 cryptic prophage gene sequences *in vivo* (García Contreras et al., 2008; Hong et al., 2010). Cryptic prophages are captured prophages that no longer produce infective phage particles but are beneficial for the stress response (Wang et al., 2010), including resuscitation from the persister state (Song et al., 2021). However, in *Escherichia coli*, cryptic prophages contain active lysis genes that must be repressed by CRISPR-Cas (Song et al., 2022).

Here, we investigate whether Hha/TomB play a role in phage defense by screening a *hha/tomB* double knockout for its sensitivity to phage infection by using the BASEL collection of 69 *Escherichia coli* lytic phages (Maffei et al., 2021). Surprisingly, we find that toxin Hha is detrimental for phage defense with 12 of 69 BASEL phages. Focusing on BASEL phage 48 (*Vequintavirus* 137.6 kb), we found Hha/TomB increases phage adsorption. In addition, we show that phages utilize Hha/TomB along with cryptic prophage lysis genes to infect *E. coli*.

## RESULTS

### BASEL phage screen with *hha/tomB*

Given that there are few studies on type VII TAs and phage inhibition (Guo et al., 2024), we chose to investigate whether type VII TAs Hha/TomB played a role in phage inhibition. Hence, we used 69 BASEL lytic phages (Humolli et al., 2025) to test the susceptibility of an *E. coli* strain lacking the *hha/tomB* locus vs. the wild-type and found 12 phages less infective with the *hha/tomB* strain at a multiplicity of infection (MOI) of 0.1: phages 31, 34, 39, 40, 41, 42, 43, 48, 50, 55, 56, and 58 (**Fig. S1**).

Of these, we tested BASEL phages 39, 41, 48, 50, 56, and 58 for adsorption changes since the *hha/tomB* deletion mutant was most resistant to these six phages (**Fig. S1**) and since changes in adsorption are one of the most frequent ways *Escherichia* spp. inhibit phage infection (Gaborieau et al., 2024). We found (**Table 2**) that BASEL phage 48 had the largest adsorption reduction among these six phages. Hence, because of its consistent and large phenotype (**Fig. S1**) and reduction in adsorption with *hha/tomB* (**Table 2**), we focused on BASEL phage 48 (**Fig. 1A**). BASEL phage 48 (henceforth ‘phage 48’) is a *Vequintavirus* (myovirus morphology) of 138 kb which attaches to the lipopolysaccharide outer core (Humolli et al., 2025). Note the *hha/tomB* and wild-type strains grew identically (**Fig. 1B**), so there were no artifacts related to differences in growth.

**Fig. 1.**
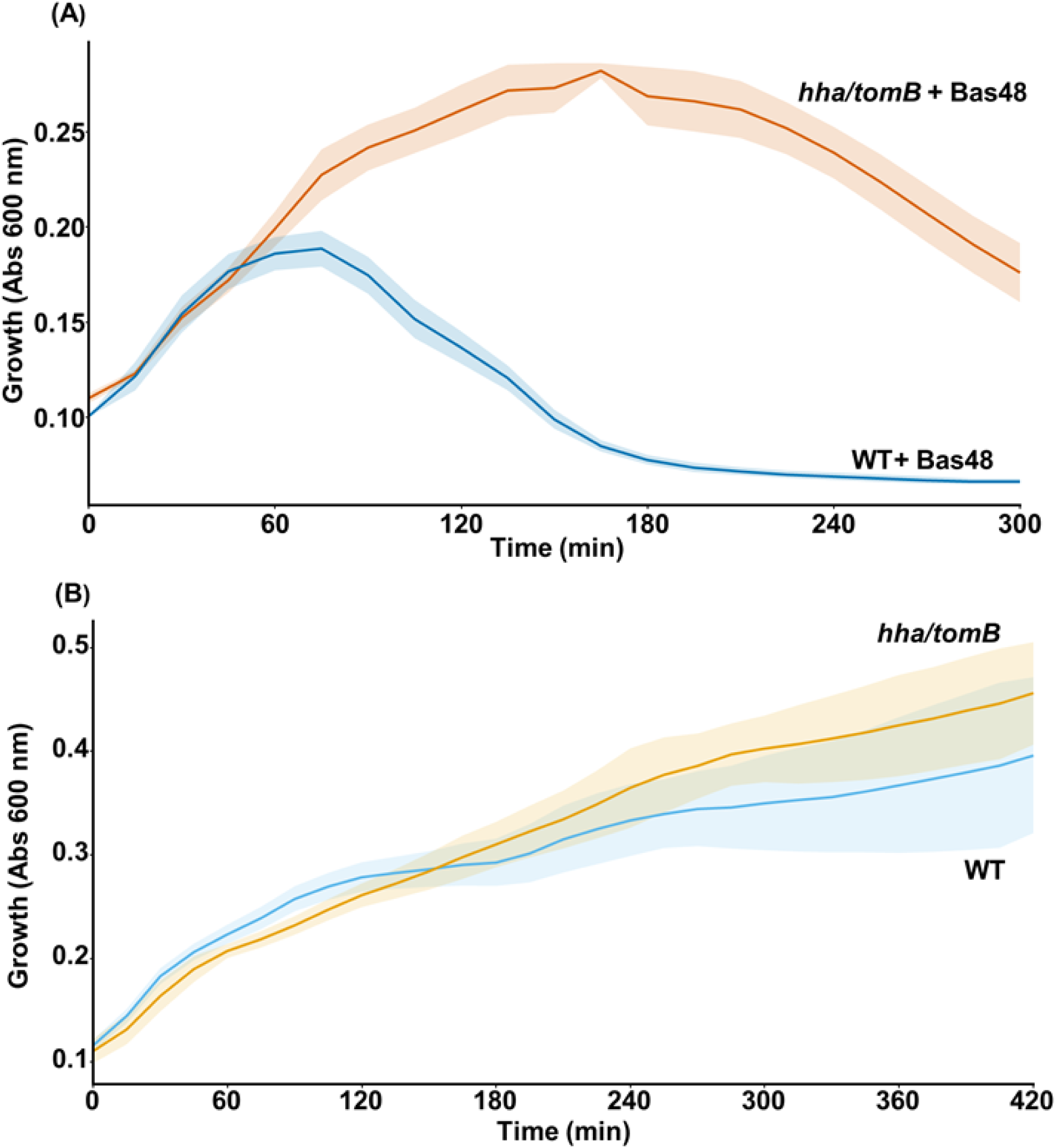
*hha/tomB* increases sensitivity to BASEL phage 48. **A**. Phages at 0.1 MOI were added to cells grown to a turbidity of 0.1 at 600 nm. Growth was recorded in a microplate Tecan reader with some shaking. Results from three independent cultures with standard deviations shown as shading. Results with BASEL phages 31, 34, 39, 40, 41, 42, 43, 48, 50, 55, 56, and 58 are shown in **Fig. S1B**. The *hha tomB* deletion does not affect growth in the absence of phage.

### Hha facilitates phage infection

To determine whether toxin Hha or antitoxin TomB was responsible for the decrease in infection with the BASEL lytic phages and the *hha/tomB* strain (**Fig. S1**), we utilized single gene deletions of *hha* and *tomB* and found that (i) the *hha* deletion reduced infection (**Fig. 2A**), whereas (ii) the *tomB* deletion increased infection (**Fig. 2A**). Hence, the two single deletion strains with phage 48 corroborated that toxin Hha is responsible for the greater sensitivity of the wild-type strain since *tomB* deletion leads to more Hha activity and this strain had the greatest sensitivity. Note there was no difference in growth for these strains in the absence of phage (**Fig. 2B**). Furthermore, the *hha* deletion was complemented with plasmid pCA24N-*hha* and 0.1 mM IPTG (**Fig. 2A**); the addition of pCA24N with 0.1 mM IPTG does not affect growth but pCA24N-*hha* with 0.1 mM IPTG reduces growth due to expression of toxin Hha, as expected (**Fig. 2B**).

**Fig. 2.**
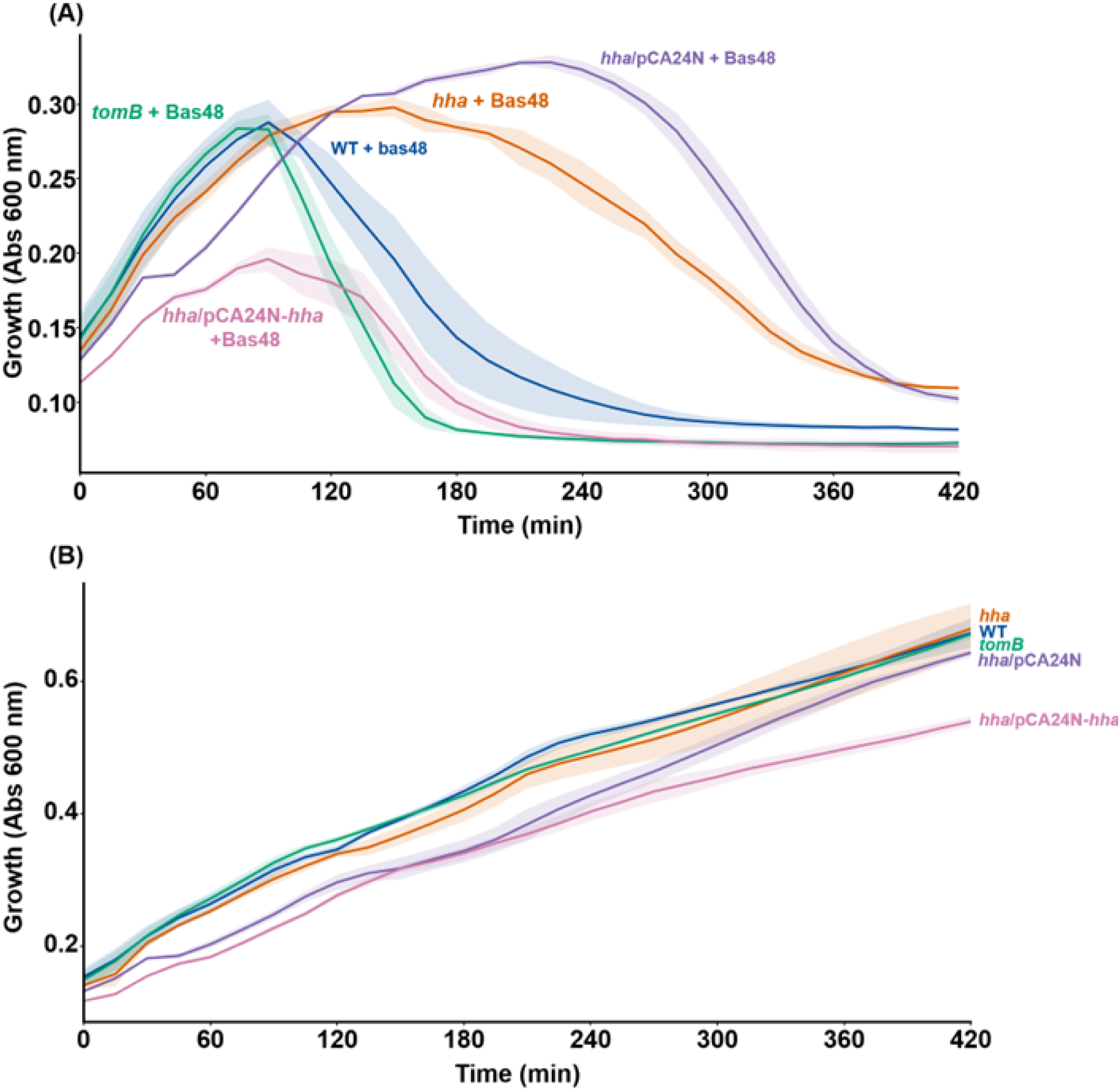
Hha increases infectivity with BASEL phage 48 and may be complemented. **A**. The *hha* deletion decreases infection, and the *tomB* deletion increases infection. BASEL phage 48 at 0.1 MOI was added to cells grown to a turbidity of 0.1 at 600 nm. The *hha* deletion was complemented with plasmid pCA24N-*hha* and 0.1 mM IPTG. **B**. The *hha* and *tomB* deletions as well as the addition of pCA24N and pCA24N-*hha* with 0.1 mM IPTG do not affect growth in the absence of phage. Results from three independent cultures with standard deviations shown as shading.

### Hha/TomB increase phage adsorption

Since the most-effective phage defense mechanism is blocking attachment (Chen et al., 2024; Gaborieau et al., 2024; David et al., 2025), we tested six of the 12 phages with the greatest inhibition of phage identified in the growth assay (**Fig S1**: 39, 41, 48, 50, 56, 58) and found that the adsorption of phage 48 was inhibited the most by the deletion of *hha/tomB*; hence, Hha/TomB increases phage adsorption 17 ± 3-fold with phage 48 (**Table 2**). Given that these six phages have a highly convergent adsorption mechanism to lipopolysaccharide (e.g., OmpC, Tsx), these results suggest a specific bacterial receptor related to membrane stress may trigger Hha activation.

### Hha functions through cryptic prophage CPS-53

Since the toxicity of Hha has been linked to increasing cell lysis and biofilm dispersal by activating cryptic prophage lytic genes *appY* and *rzpD* of DLP12 and *alpA* and *yfjZ* of CP4-57 by inducing protease ClpXP (García Contreras et al., 2008) and since Hha induces excision of prophages CP4-57 and DLP12 of *E. coli* by inducing *intA* and *alpA* (Wang et al., 2009), we investigated if the increase in sensitivity of *E. coli* to phage 48 was related to its nine cryptic prophages (Wang et al., 2010) (**Fig. S2)**. Hence, we measured the ability of phage 48 to infect cells with each of the cryptic prophages deleted (Wang et al., 2010) and found that only deleting cryptic prophage CPS-53 resulted in increased cell survival compared to the wild-type strain (**Fig. 3A**), whereas deletion of the other eight cryptic prophages resulted in slight to moderate decreases in cell survival (**Fig. S2**). In the absence of phage, there was no difference in growth (**Fig. 3B**). Therefore, Hha functions through cryptic prophage CPS-53 to increase infection with phage 48.

**Fig. 3.**
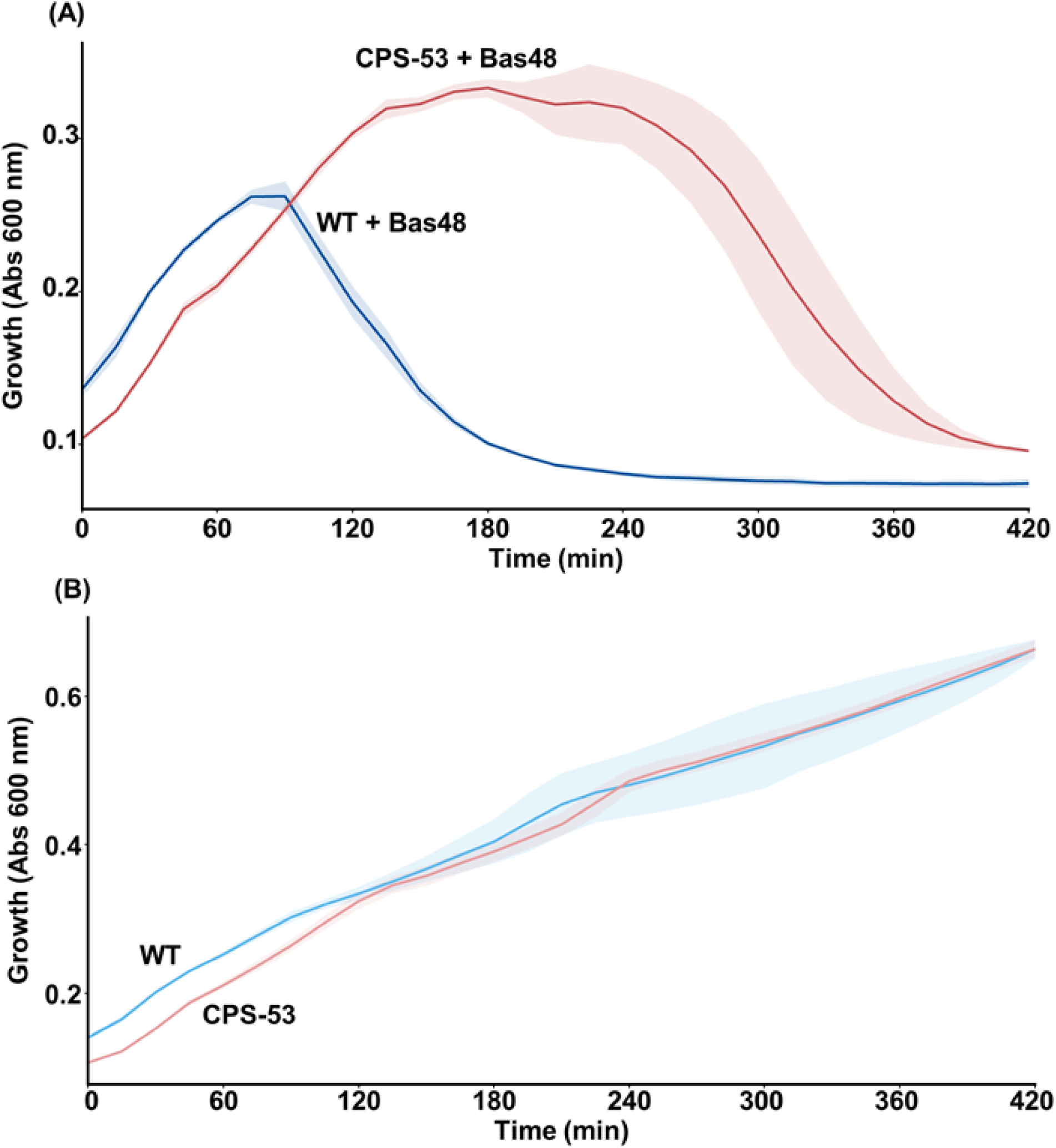
Cryptic prophage CPS-53 increases infectivity with BASEL phage 48. **A**. Deletion of cryptic prophage CPS-53 (ΔCPS-53) decreases infection at MOI 0.1. **B**. The ΔCPS-53 deletion does not affect growth in the absence of phage. Results from three independent cultures with standard deviations shown as shading. Results with all nine cryptic prophages are shown in **Fig. S2**.

### Hha functions through lytic genes in cryptic prophages

To explore further the role of CPS-53 and phage 48, we investigated all ten single gene mutations in cryptic prophage CPS-53 and found *yfdH* (encodes glucosyl transferase) and *yfdO* (encodes a protein that enhances the host response to stress) were less sensitive to phage 48 (**Fig. 4A**). Hence, phage 48 relies on cryptic prophage proteins YfdH and YfdO. Since phage 48 relies on CPS-53 proteins YfdH and YfdO, we hypothesized that phage 48 may also rely on lysis genes from other cryptic prophages, since Hha activates cryptic prophage lytic genes (García Contreras et al., 2008; Hong et al., 2010). Therefore, we investigated the seven lysis-related genes of cryptic prophages DLP12 (*essD, ybcS, rzpD*, and *rzoD*) and Qin (*ydfQ, essQ,* and *ydfD*) since Hha was previously found to bind within DLP12 and Qin (García Contreras et al., 2008). Remarkably, we found deleting all but *rzpD* of these cryptic prophage lysis genes rendered the *E. coli* host less sensitive to phage 48 (**Fig. 4A**) without affecting growth (**Fig. 4B**), with the largest effect for deleting *essQ* and *ydfD* from Qin and *essD, ybcS,* and *rzoD* from DLP12. Of these, *rzoD* and *essD* had the greatest impact of reducing phage 48 infection based on the percentage of OD obtained in the absence of phage (**Fig. 4C**). Corroborating these results for three of the deletions with the most-prevalent phenotypes, the *essD, essQ,* and *yfdH* deletions could be complemented (**Fig. 5**). Hence, phage 48, via Hha, uses the cryptic prophage proteins, including the lysis genes, to infect the cell.

**Fig. 4.**
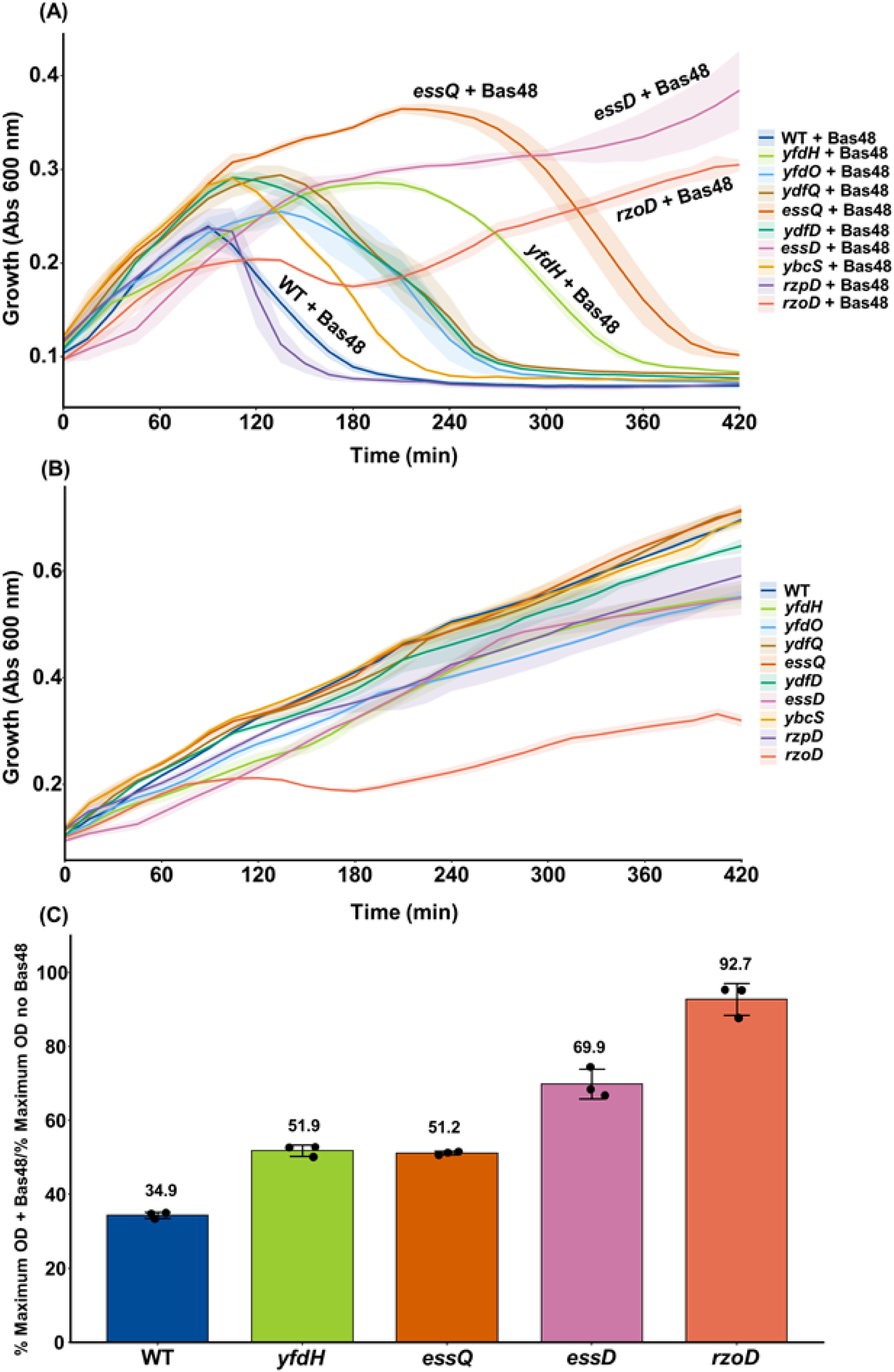
Cryptic prophage lysis proteins increase infectivity with BASEL phage 48. **A**. Deletion of cryptic prophage lysis genes from Qin and DLP12 decreases infection at MOI 0.1 along with several deletions in cryptic prophage CPS-53 (*yfdH* and *yfdO*). **B**. The deletion of lysis genes does not affect growth in the absence of phage. Results from three independent cultures with standard deviations shown as shading. Results with all single gene deletions of CPS-53are shown in **Fig. S3**.

**Fig. 5.**
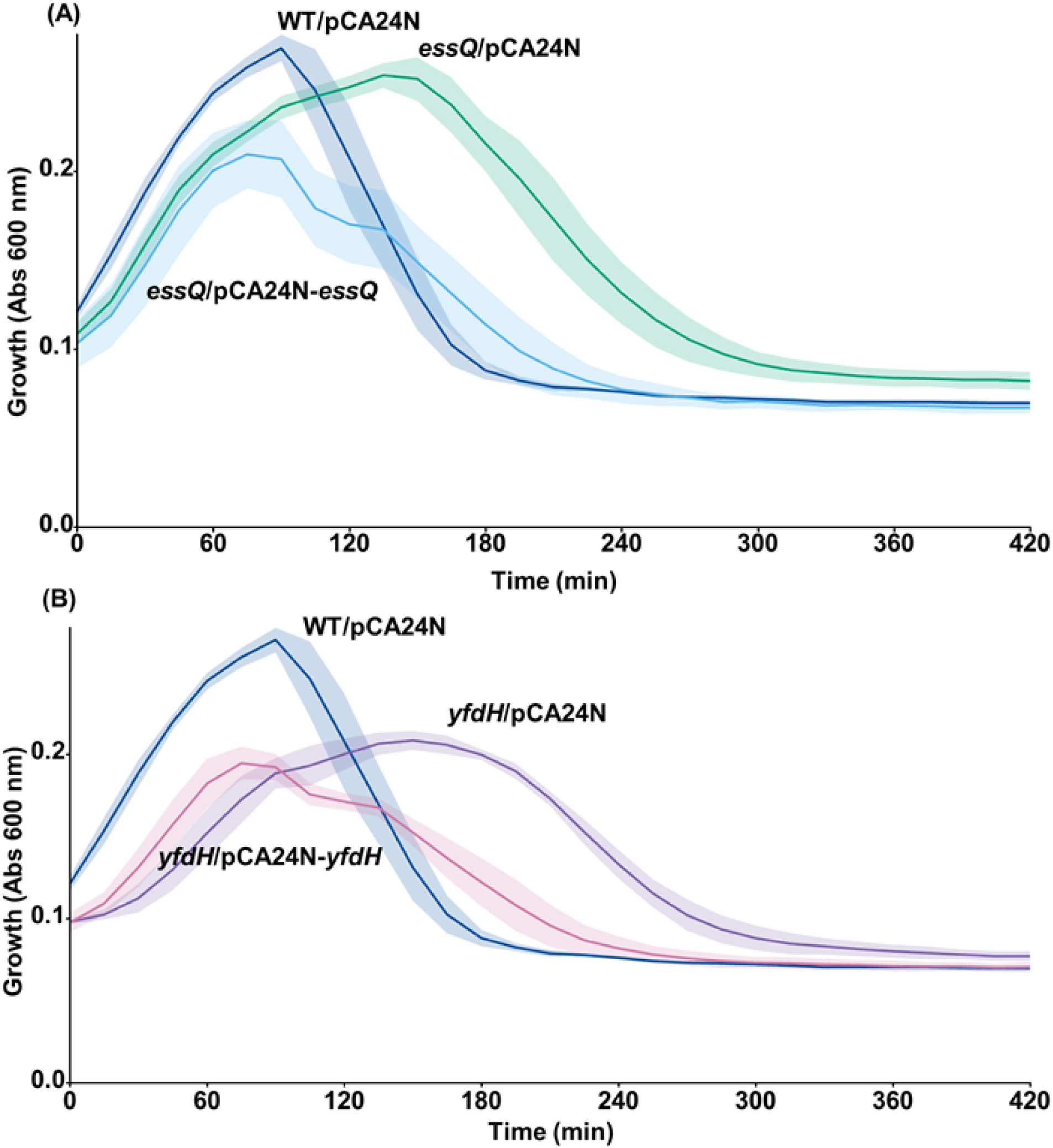
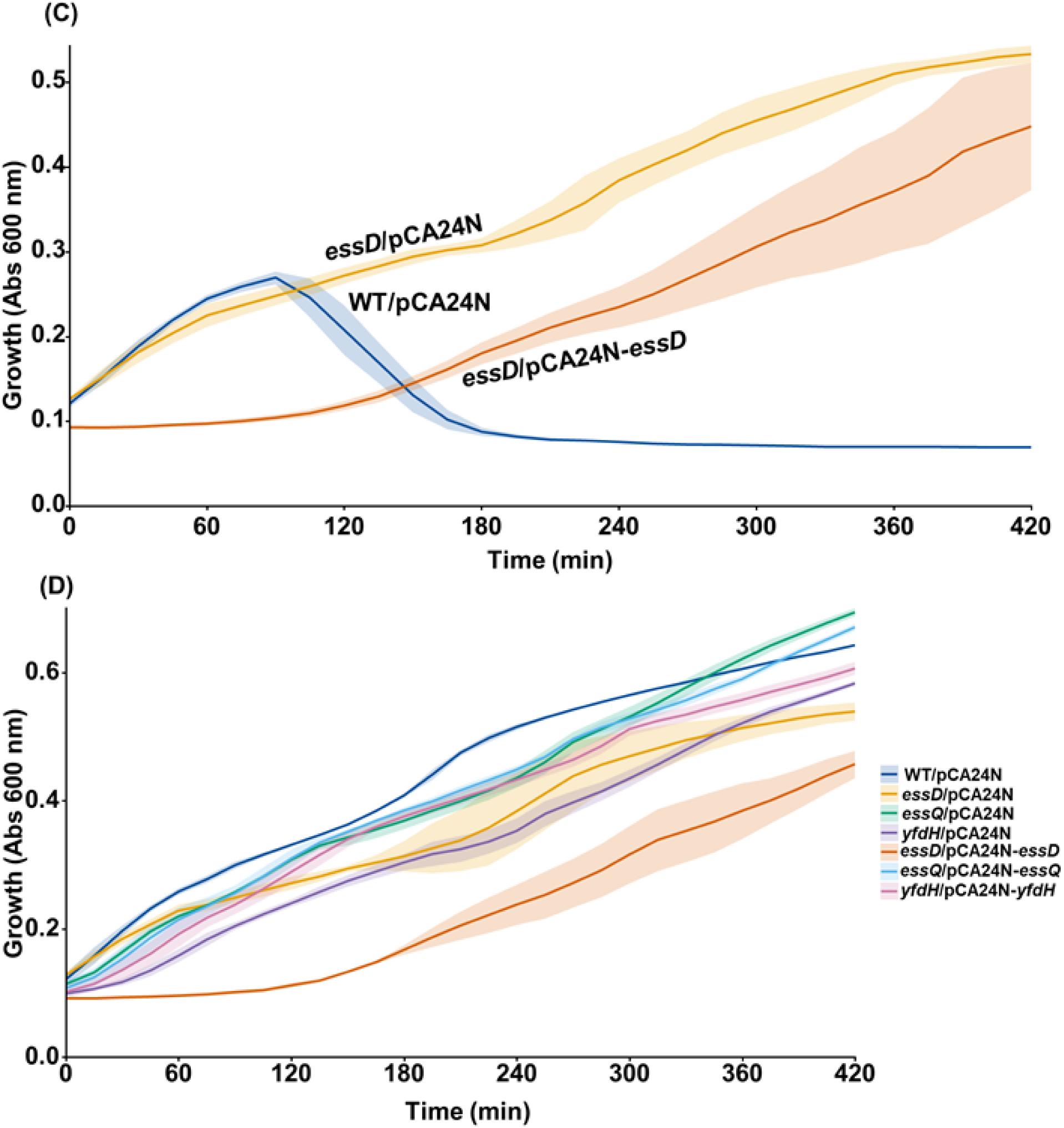
Deletion of cryptic prophage lysis proteins EssQ (Qin) and EssD (DLP12) and glucosyl transferase YfdH (CPS-53) may be complemented with BASEL phage 48. **A**. Complementation of *essQ* (encodes lysis protein from Qin cryptic prophage) deletion. B. Complementation of *essD* (encodes lysis protein from DLP12 cryptic prophage) deletion. C. Complementation of *yfdH* (encodes putative glucosyl transferase from CPS-53 cryptic prophage) deletion. D. Growth in the absence of phage. Results from three independent cultures with MOI 0.1 with standard deviations shown as shading.

## DISCUSSION

In this report, we show clearly the surprising result that the chromosomally-encoded toxin Hha of the Hha/TomB TAs is detrimental for phage defense for 12 of the 69 BASEL collection of lytic phages; i.e., infection by these 12 phages is enhanced through Hha. This result is contrary to the current paradigm (Song and Wood, 2020) that TAs inhibit phage infection. Moreover, by linking Hha to cryptic prophages and by studying single-gene deletions in the cryptic prophages, we determined the mechanism for the reduction in sensitivity with phage 48 is that Hha is utilized by phage 48 to induce the lytic genes of the *E. coli* cryptic prophages; i.e., phage 48 utilizes Hha to lyse the cell by making use of the ancient lysis genes of cryptic prophages captured by *E. coli* (**Fig. 6**).

**Fig. 6.**
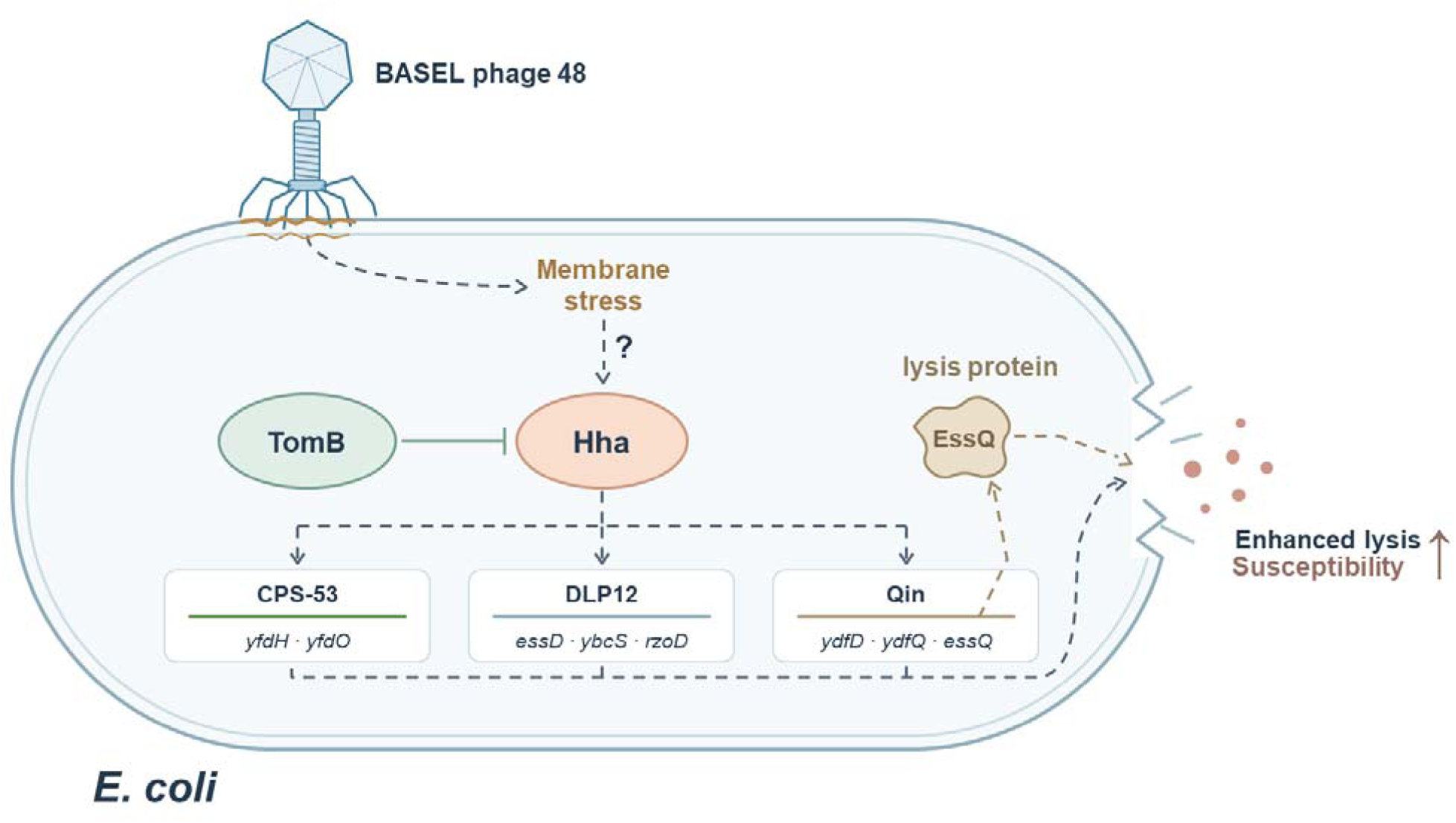
Schematic of Hha activation by BASEL phage 48. Upon adsorption, BASEL phage 48 induces membrane stress, which activates Hha of the Hha/TomB toxin/antitoxin system through an unknown mechanism. Activated Hha then induces CPS-53, DLP-12, and Qin cryptic prophage genes, some of which produce cell lysis proteins and facilitate phage infection (e.g., EssQ).

This novel mechanism fits well with an earlier result that the TAs Doc/Phd of prophage P1 requires the chromosomal *E. coli* TAs MazF/MazE to inhibit protein synthesis. Hence, prophages can depend on chromosomal TAs (Hazan et al., 2001) to control cell physiology. Similarly, we now show lytic phages can depend on cryptic prophages to lyse the cell via TAs; i.e., phages can *manipulate* the lysis genes of cryptic prophages through toxin Hha (**Fig. 6**). This is in opposition to the paradigm that prophages primarily protect the host (Azulay et al., 2026).

Since H-NS inhibits partially *E. coli* CRSPR-Cas (Shmakov et al., 2017) and since we found CRISPR-Cas inhibits *E. coli* cryptic prophage functional lytic genes such as *ydfD* of Qin and *rzoD* of DLP12 (Song et al., 2022), it is reasonable that Hha, a partner of H-NS, regulates the expression of other cryptic prophage lytic proteins, such as and *ydfD* and *rzoD* as we found here.

Although we focused on the role of Hha of the Hha/TomB TA system and cryptic prophage lysis genes including *ydfQ, essQ,* and *ydfD* from the Qin cryptic prophage and *essD, ybcS, rzpD*, and *rzoD* from the DLP12 cryptic prophage, Hha is pleiotropic since its deletion affects up to 6% of the genome in biofilms (García Contreras et al., 2008). Moreover, using nickel enrichment microarrays, we found that Hha binds at least 70 genomic regions, including 15 areas within the DLP12, CP4-57, and Qin cryptic prophage regions (García Contreras et al., 2008) as well as controls excision of cryptic prophages CP4-57 and DLP12 (Wang et al., 2009). Hence, the discovery that Hha influences BASEL group phages via the CPS-53 and Qin cryptic prophage-related proteins is consistent, but other mechanisms may be possible due to Hha influencing other sites in the *E. coli* genome and due to its partnership with H-NS.

In addition, it seems deleterious for *E. coli* to harbor the *hha/tomB* locus since Hha may be utilized by lytic phages like phage 48 to increase infection. However, the physiological role of Hha may be instead to induce persistence during lytic phage infection, as was found recently for MqsR of the MqsR/MqsA/MqsC TAs with T2 phage (Fernández-García et al., 2024); hence, the host may be willing to sacrifice some viable cells by activating toxin Hha during phage infection while increasing persistence to ensure that some cells survive infection. Or perhaps some lytic phages such as phage 48 have learned to manipulate the tools used to silence foreign DNA (H-NS and Hha), which have become an Achilles’s heel, the irony being that tools that silence foreign DNA may actually facilitate foreign invaders.

## METHODS

### Bacterial strains, phages, and growth conditions

The bacterial strains and isogenic mutants and plasmids are described in **Table 1**. The BASEL collection of *E. coli* lytic phages (Maffei et al., 2021) was provided by Alexander Harms. Cells were grown in lysogeny broth (LB, 1 % tryptone, 0.5 % yeast, and 1 % NaCl w/v) at 37°C at 250 rpm. Kanamycin (50 µg/mL) was used for preculturing knockout mutants where applicable, and chloramphenicol (30 µg/mL) was added to strains containing pCA24N-based plasmids to maintain the vector.

**Table 1.**
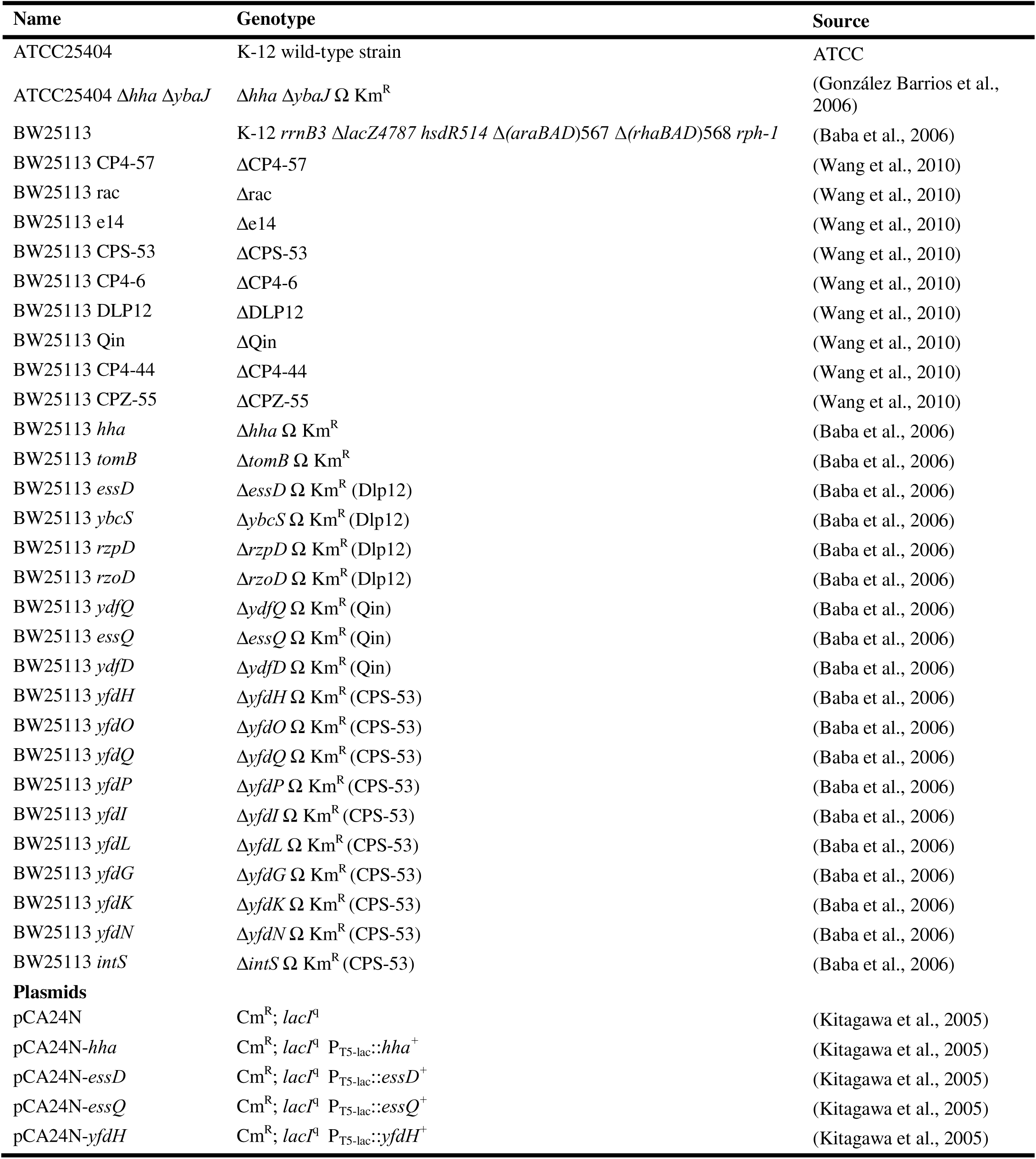
*Escherichia coli* strains and plasmids used in this study. Amp^R^, ampicillin resistance; Km^R^, kanamycin resistance.

**Table 2.** Adsorption (%) of Basel phage to *E. coli* strains after 8 min of incubation (n = 3 independent cultures). Values are presented as mean ± standard deviation (stdev, %).

| Basel phage | WT (%) | | | mean ± stdev (%) | $\Delta hha \Delta tomB$ (%) | | | mean ± stdev (%) | Fold change* ($\Delta hha \Delta tomB$/WT) |
| --- | --- | --- | --- | --- | --- | --- | --- | --- | --- |
| 39 | 7 | 4 | 1 | 4 ± 3 | 2 | 2 | 3 | 2.3 ± 0.6 | -1.7-fold |
| 41 | 0.7 | 0.9 | 0.5 | 0.7 ± 0.2 | 0.1 | 0.1 | 0.1 | 0.1 ± 0 | -7-fold |
| **48** | 50 | 60 | 60 | 56.7 ± 5.8 | 3 | 4 | 3 | 3.3 ± 0.6 | **-17-fold** |
| 50 | 0.8 | 0.9 | 1.5 | 1.1 ± 0.4 | 14 | 11 | 13 | 12.7 ± 1.5 | +12-fold |
| 56 | 30 | 30 | 40 | 33.3 ± 5.8 | 23 | 28 | 21 | 24 ± 4 | -1.4-fold |
| 58 | 90 | 100 | 85 | 91.7 ± 7.6 | 40 | 40 | 30 | 36.7 ± 5.8 | -2.5-fold |

\* Negative (–) and positive (+) signs indicate decreased and increased phage adsorption, respectively, in $\Delta hha \Delta tomB$ relative to WT.

### Temporal phage assay

Cultures were grown overnight in LB with antibiotics, as appropriate, diluted 1:100 into LB with no antibiotics, incubated to a turbidity of around 0.1 at 600 nm, and transferred to a 96-well plate and Basel phages were then added at MOI 0.1. Growth was recorded in the microplate Tecan reader with periodic shaking.

### Adsorption assay

Phage adsorption was performed as described previously (Fernández-García et al., 2024) by diluting overnight LB cultures 1:100 and incubating until turbidity (600 nm) ∼ 0.5, then phage was added at MOI 0.01 and incubated at room temperature without shaking for 8 min. Samples (1 mL) were centrifuged for 10 min at 5,000 rpm, rinsed once with phosphate-buffered saline (PBS, 8 g NaC1, 0.2 g KC1, 1.15 g Na_2_HPO_4_, and 0.2 g KH_2_PO_4_ in ddH_2_O 1000 mL) and vortexed 15 sec at maximum (Genie 2, Fisher scientific.) to remove free and loosely-attached phages and the cells were resuspended in phage buffer (0.1 M NaCl, 10 mM MgSO_4_, and 20 mM Tris-HCl pH 7.5), and PFU were determined using double layer agar plates.

## ACKNOWLEDGEMENTS

This work was supported by funds derived from NSF grant 2515448 (UEI: NPM2J7MSCF61). We thank Alexander Harms for providing the BASEL collection of *Escherichia coli* phages.

## Supplemental Information

**Fig. S1.**
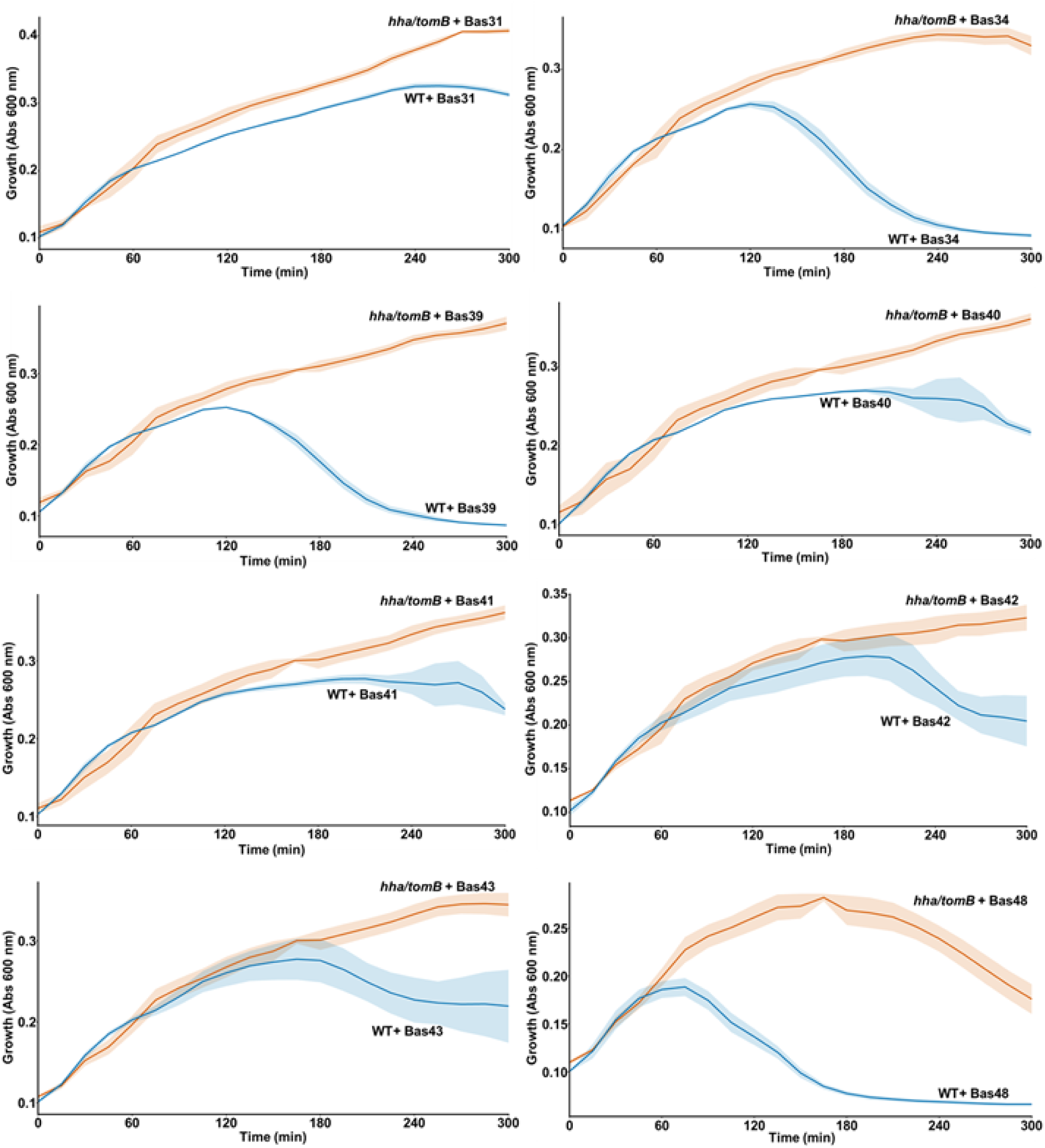
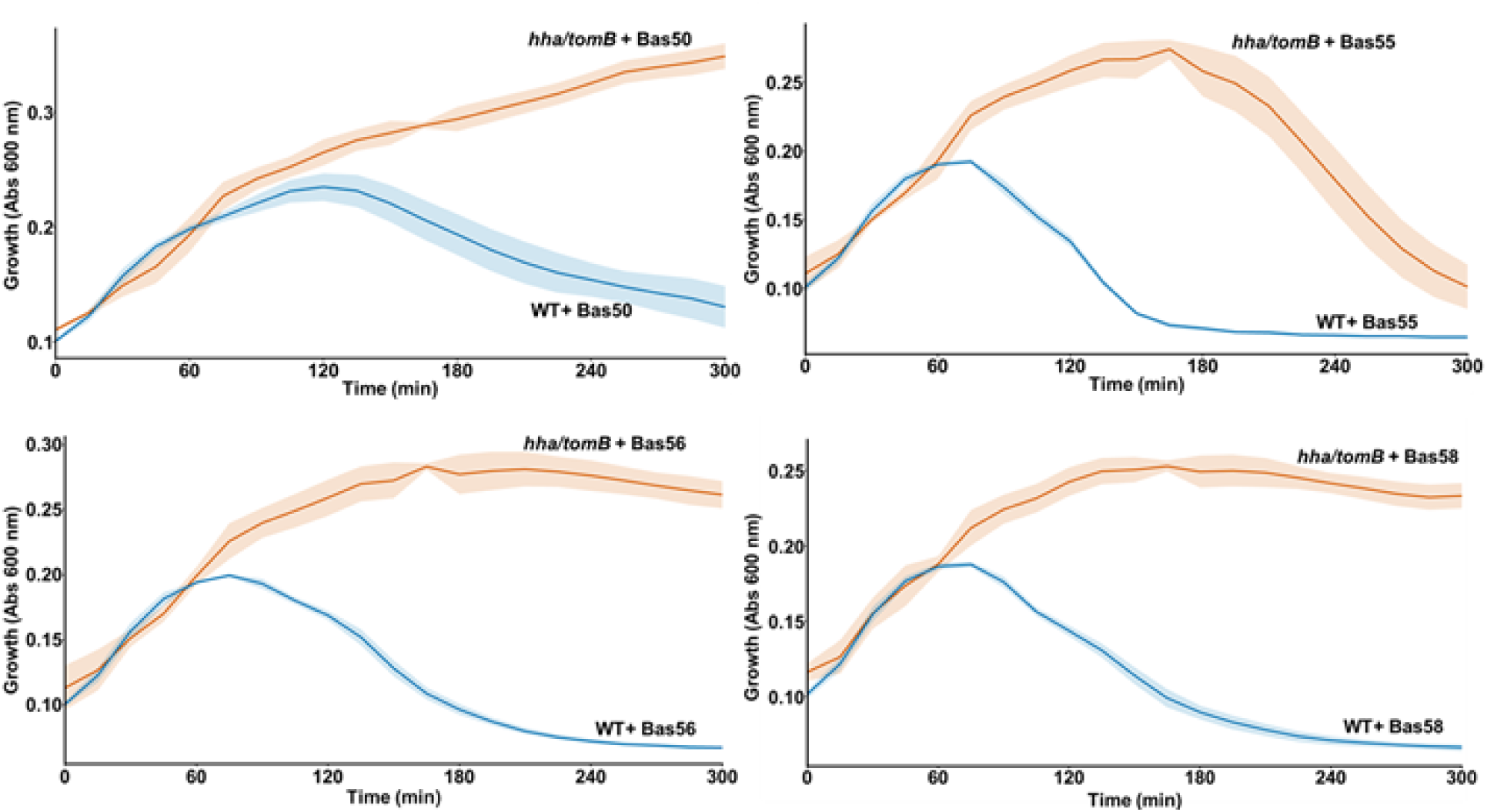
*Hha/tomB* inhibits BASEL phages 31, 34, 39, 40, 41, 42, 43, 48, 50, 55, 56, and 58. Three independent cultures with standard deviations shown as shading and MOI 0.1.

**Fig. S2.**
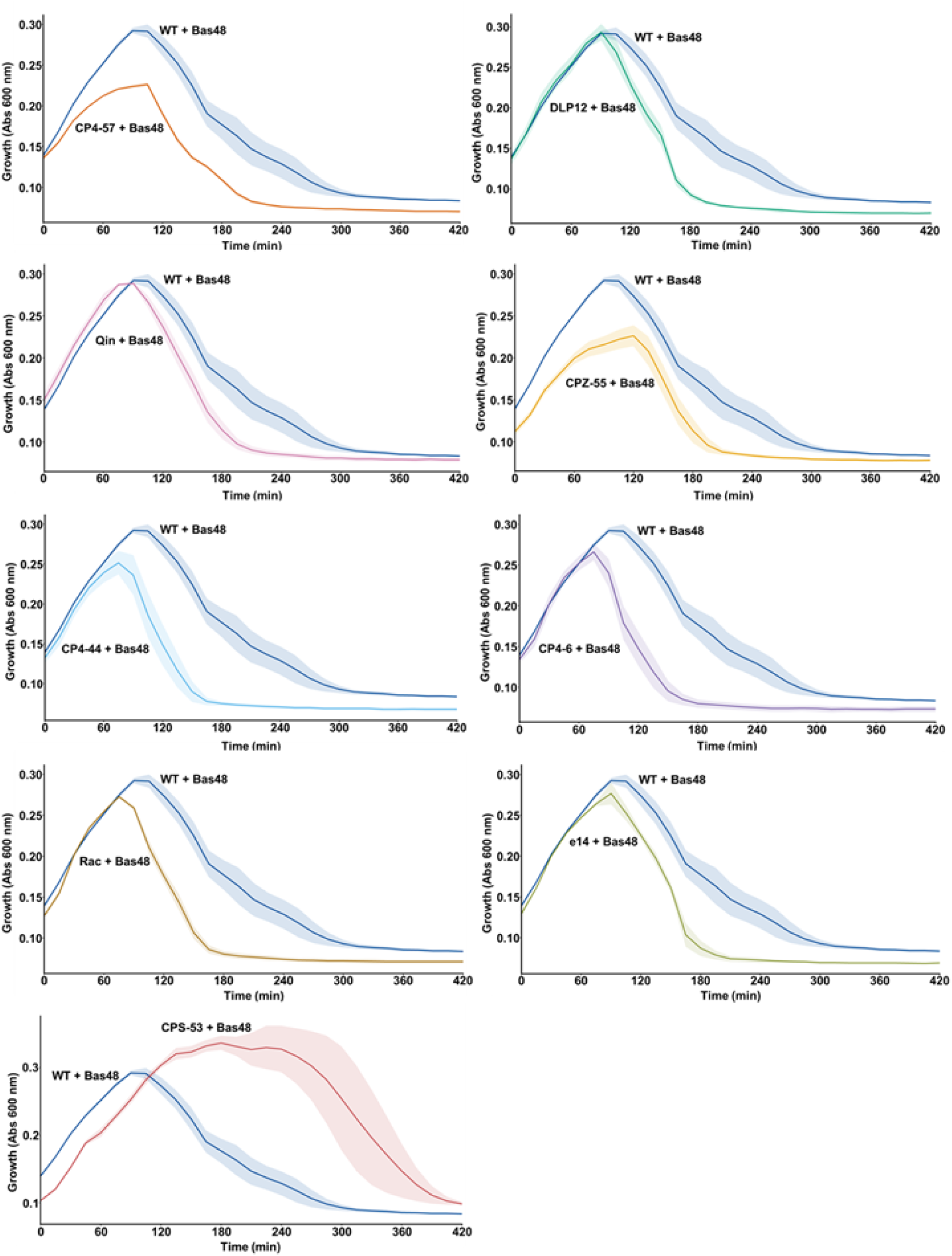
Nine delta cryptic prophage strains (including CPS-53) and BASEL phage 48. Three independent cultures with standard deviations shown as shading and MOI 0.1.

**Fig. S3.**
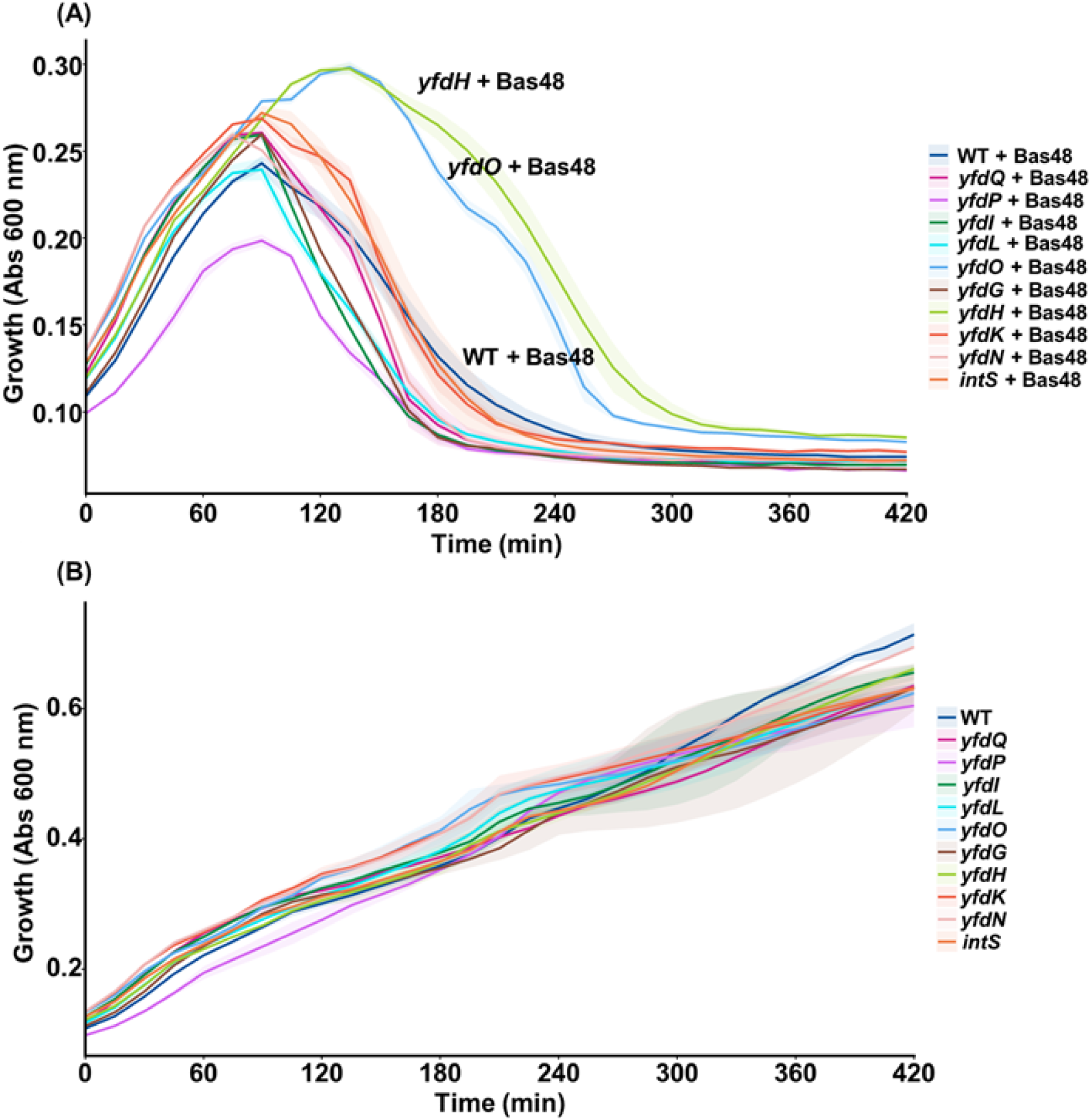
Screen for CPS-53 single gene deletions and BASEL phage 48. One independent culture with standard deviations shown for the two technical replicates as shading and MOI 0.1.

## REFERENCES

Azulay, G., Sigal, N., Tzohar Rabinovich, K., Cohen, Y., Das, A., Koenigsfeld, D. et al. (2026) A prophage-encoded anti-phage defence system that prevents DNA packaging by targeting the terminase complex. Nature Communications.

Baba, T., Ara, T., Hasegawa, M., Takai, Y., Okumura, Y., Baba, M. et al. (2006) Construction of *Escherichia coli* K-12 in-frame, single-gene knockout mutants: the Keio collection. Mol Syst Biol 2: 2006 0008.

Chen, L., Zhao, X., Wongso, S., Lin, Z., and Wang, S. (2024) Trade-offs between receptor modification and fitness drive host-bacteriophage co-evolution leading to phage extinction or co-existence. The ISME Journal 18.

David, E., Plantady, C., Poissonnier, S., Elliott, J.F.K., Kenck, E., Le Boulch, J. et al. (2025) Systematic functional assessment of anti-phage systems in their native host. Philosophical Transactions of the Royal Society B: Biological Sciences 380.

Fernández-García, L., Song, S., Kirigo, J., Battisti, M.E., Petersen, M.E., Tomás, M., and Wood, T.K. (2024) Toxin/antitoxin systems induce persistence and work in concert with restriction/modification systems to inhibit phage. Microbiology Spectrum 12: e03388–03323.

Fineran, P.C., Blower, T.R., Foulds, I.J., Humphreys, D.P., Lilley, K.S., and Salmond, G.P.C. (2009) The phage abortive infection system, ToxIN, functions as a protein–RNA toxin–antitoxin pair. Proceedings of the National Academy of Sciences 106: 894–899.

Gaborieau, B., Vaysset, H., Tesson, F., Charachon, I., Dib, N., Bernier, J. et al. (2024) Prediction of strain level phage–host interactions across the Escherichia genus using only genomic information. Nature Microbiology 9: 2847–2861.

García Contreras, R., Zhang, X.-S., Kim, Y., and Wood, T.K. (2008) Protein Translation and Cell Death: The Role of Rare tRNAs in Biofilm Formation and in Activating Dormant Phage Killer Genes. PLoS ONE 3: e2394.

González Barrios, A.F., Zuo, R., Ren, D., and Wood, T.K. (2006) Hha, YbaJ, and OmpA Regulate *Escherichia coli* K12 Biofilm Formation and Conjugation Plasmids Abolish Motility. Biotechnology and Bioengineering 93: 188–200.

Guo, Y., Tang, K., Sit, B., Gu, J., Chen, R., Shao, X. et al. (2024) Control of lysogeny and antiphage defense by a prophage-encoded kinase-phosphatase module. Nature Communications 15: 7244.

Hazan, R., and Engelberg-Kulka, H. (2004) *Escherichia coli mazEF*-mediated cell death as a defense mechanism that inhibits the spread of phage P1. Mol Genet Genomics 272: 227–234.

Hazan, R., Sat, B., Reches, M., and Engelberg-Kulka, H. (2001) Postsegregational Killing Mediated by the P1 Phage “Addiction Module” *phd-doc* Requires the *Escherichia coli* Programmed Cell Death System *mazEF*. Journal of Bacteriology 183: 2046–2050.

Hong, S.H., Lee, J., and Wood, T.K. (2010) Engineering global regulator Hha of *Escherichia coli* to control biofilm dispersal. Microb Biotechnol 3: 717–728.

Humolli, D., Piel, D., Maffei, E., Heyer, Y., Agustoni, E., Shaidullina, A. et al. (2025) Completing the BASEL phage collection to unlock hidden diversity for systematic exploration of phage–host interactions. PLOS Biology 23: e3003063.

Jurėnas, D., Fraikin, N., Goormaghtigh, F., and Van Melderen, L. (2022) Biology and evolution of bacterial toxin–antitoxin systems. Nature Reviews Microbiology 20: 335–350.

Kitagawa, M., Ara, T., Arifuzzaman, M., Ioka-Nakamichi, T., Inamoto, E., Toyonaga, H., and Mori, H. (2005) Complete set of ORF clones of *Escherichia coli* ASKA library (a complete set of *E. coli* K-12 ORF archive): unique resources for biological research. DNA Res 12: 291–299.

Maffei, E., Shaidullina, A., Burkolter, M., Heyer, Y., Estermann, F., Druelle, V. et al. (2021) Systematic exploration of Escherichia coli phage–host interactions with the BASEL phage collection. PLOS Biology 19: e3001424.

Marimon, O., Teixeira, J.M.C., Cordeiro, T.N., Soo, V.W.C., Wood, T.L., Mayzel, M. et al. (2016) An oxygen-sensitive toxin–antitoxin system. Nature Communications 7: 13634.

Pecota, D.C., and Wood, T.K. (1996) Exclusion of T4 Phage by the *hok/sok* Killer Locus from Plasmid R1. J Bacteriol 178: 2044–2050.

Pu, H., Chen, Y., Zhao, X., Dai, L., Tong, A., Tang, D. et al. (2025) A toxin-antitoxin system provides phage defense via DNA damage and repair. Nature Communications 16: 3141.

Ren, D., Bedzyk, L.A., Thomas, S.M., Ye, R.W., and Wood, T.K. (2004) Gene expression in *Escherichia coli* biofilms. Appl Microbiol Biotechnol 64: 515–524.

Rimsky, S. (2004) Structure of the histone-like protein H-NS and its role in regulation and genome superstructure. Curr Opin Microbiol 7: 109–114.

Seo, E., Hwang, H.-J., Kim, Y., and Wood, T.K. (2026) Type IX class utilizing cell signals for organizing toxin-antitoxin systems. Trends in Microbiology 34: 735.

Shmakov, S.A., Sitnik, V., Makarova, K.S., Wolf, Y.I., Severinov, K.V., and Koonin, E.V. (2017) The CRISPR Spacer Space Is Dominated by Sequences from Species-Specific Mobilomes. mBio 8: e01397–01317.

Song, S., and Wood, T.K. (2020) A Primary Physiological Role of Toxin/Antitoxin Systems Is Phage Inhibition. Front Microbiol 11: 1895.

Song, S., Kim, J.-S., Yamasaki, R., Oh, S., Benedik, M.J., and Wood, T.K. (2021) *Escherichia coli* cryptic prophages sense nutrients to influence persister cell resuscitation. Environ Microbiol 23: 7245–7254.

Song, S., Semenova, E., Severinov, K., Fernández-García, L., Benedik, M.J., Maeda, T., and Wood, T.K. (2022) CRISPR-Cas Controls Cryptic Prophages. International Journal of Molecular Sciences 23: 16195.

Wang, X., Kim, Y., and Wood, T.K. (2009) Control and Benefits of CP4-57 Prophage Excision in *Escherichia coli* Biofilms. ISME J 3: 1164–1179.

Wang, X., Yao, J., Sun, Y.-C., and Wood, T.K. (2021) Type VII Toxin/Antitoxin Classification System for Antitoxins that Enzymatically Neutralize Toxins. Trends in Microbiology 29: 388–393.

Wang, X., Kim, Y., Ma, Q., Hong, S.H., Pokusaeva, K., Sturino, J.M., and Wood, T.K. (2010) Cryptic prophages help bacteria cope with adverse environments. Nat Commun 1: 147.

Yamaguchi, S., Fernandez, S.G., Wassarman, D.R., Lüders, M., Schwede, F., and Kranzusch, P.J. (2025) Activating and inhibiting nucleotide signals coordinate bacterial anti-phage defense. bioRxiv: 2025.2007.2009.663793.

